# srnaMapper: an optimal mapping tool for sRNA-Seq reads

**DOI:** 10.1101/2021.01.12.426326

**Authors:** Matthias Zytnicki, Christine Gaspin

## Abstract

**Motivation:** Sequencing is the key method to study the impact of short RNAs, which include micro RNAs, tRNA-derived RNAs, and piwi-interacting RNA, among other. The first step to make use of these reads is to map them to a genome. Existing mapping tools have been developed for the long RNAs in mind, and, so far, no tool has been conceived for short RNAs. However, short RNAs have several distinctive features which make them different from messenger RNAs: they are shorter (not greater than 200bp), they often redundant, they can be produced by duplicated *loci*, and they may be edited at their ends.

**Results:** In this work, we present a new tool, srnaMapper, that maps these reads with all these objectives in mind. We show on two data sets that srnaMapper is more efficient considering computation time and edition error handling: it quickly retrieves all the hits, with arbitrary number of errors.

**Availability:** srnaMapper source code is available at https://github.com/mzytnicki/srnaMapper.

## 1 Introduction

Eukaryotic small RNAs (sRNAs) are defined as <200-bp long, usually untranslated, RNAs. They have been shown to participate in many aspects of cell life [1, 2].

They are generally classified according to their specific size range, biogenesis, and functional pathway. Among them, microRNAs (miRNAs) are certainly the most studied, but many other small RNAs have been shown to have a key role in regulation: tRNA-derived small RNAs (tsRNAs), small interfering RNAs (siRNAs), piwi-associated RNAs (piRNAs) and repeat-associated siRNAs (rasiRNAs), to name a few.

After the sequencing, the first task is usually to map the reads to the genome, *i.e*. predict the putative *loci* which may have produced the reads. Many mapping tools have been created so far, but none has been developed especially for sRNAs. User then resort to DNA mapping tools such as bowtie (Langmead *et al*., 2009), bowtie2 (Langmead and Salzberg, 2012), or bwa (Li and Durbin, 2009), with tuned parameters. Downstream tools may then be applied to filter the results.

Here, we present a new tool for efficiently mapping sRNA reads. It addresses all the particularities of these reads efficiently.

First, sRNA-producing *loci* are often duplicated: miRNAs are sometimes grouped into families, which generate highly similar or identical RNAs, and piRNAs are produced in interaction with transposable elements, which are known to be duplicated. Our tools provides all the hits (up to user given threshold) for each read.

Second, some sRNAs, such as miRNAs, undergo editing at their ends. Both the 5’ or the 3’ can be shrinked, extended with a template, or both. Contrary to other tools, such as bwa or bowtie, we did not set any “seed” at the ends of the reads. Moreover, the tool requires a maximum number of errors (which can be mismatches or indels), and not a percentage, since the editing is, as far as we know, not dependent of the size of the read.

Third, sRNAs are short, usually < 30bp long, and their are highly redundant (the same sRNA may be sequenced thousands times). As a result, we can store all the reads in a tree, which fits into memory, and substantially accelerate the mapping process, since the same sRNA is mapped only once, even though it is sequenced several times.

Last, our experience in sRNA-Seq showed us that the users usually want all the hits that map with the lowest number of errors. These feature is usually implemented with the option `--best --strata` in bowtie1, but is not available in every mapping tool.

## 2 Methods

First, a method based on *q*-gram filtering cannot used here, since 21bp long miRNAs, with 2 errors, should be split into *q*-grams that are too short to be useful.

In our implementation, the genome is indexed using the bwa suite, which creates a suffix array, together with the BW transform and the FM index. Since we will manipulate this structure like a tree, for the clarity of the discussion, we will refer to this structure as the genome tree, even though it is, *stricto sensu*, an array. The tools then stores the reads into a radix tree, where each path from the root to a terminal node stores a sequence. The number of reads for the corresponding sequence, as well as the read quality, is also stored into the terminal node.

Given a threshold *k* and a terminal node in the reads tree, the aim is then to find all the nodes in the genome tree with the minimal edit distance, not greater than *k* (when they exist). If *k* = 0, the problem reduces to finding the common sub-tree of the genome tree and the reads tree. If *k* > 1, the problem could be described as an “approximate” subtree search. To the best of our knowledge, the latter problem has never been described so far.

To map the reads, we first map the reads root node to the genome tree with at most *k* errors. We thus have a list of corresponding genome nodes. Then, we add a nucleotide from the reads tree: we try the new corresponding genome nodes using the previously computed list. We recursively traverse the reads tree this way to find all the matching genome nodes, and report the results when we find a terminal node.

We implemented several optimizations. First, we do not store one genome tree, but 4^8^ trees, which start with all the 8-mers. This saves time and space, since, in our data, virtually all the 8-mers were seen in the first 8bp of the reads. Second, we first try to map a read with 0 error. If it fails, the search is backtracked to the last point where a mapping with 1 error was done, etc. Third, almost the search (except loading the genome tree) can performed in parallel. Fourth, several fastq files can be given to the tool, and they will be mapped simultaneously. We tried other optimizations, but they did not given significant improvements.

## 3 Results

We compared our approach with several different tools. First, we use the widely used tool bowtie (Langmead *et al*., 2009), bowtie2 (Langmead and Salzberg, 2012), and bwa (Li and Durbin, 2009), with the parameters suggested by the review (Ziemann *et al*., 2016). We also tried several “all mapper”, such as Yara (Siragusa *et al*., 2013), which are tools designed to quickly retrieve all hits. Note that Yara does not make it possible to specify a fixed edit distance. Instead, the user can specify an error rate, which is the percentage of errors, given the read size. We choose an error rate of 10 (which is two errors at most for a read or size 20), and discarded reads with more that 2 erros. Other all mappers, such as FEM (Zhang *et al*., 2018), Hobbes (Ahmadi *et al*., 2011), and BitMapper2 (Cheng *et al*., 2019), could not be used, because the reads were too short for an edit distance of 2.

Results on the two datasets show that srnaMapper mapps all the reads that the other map (Fig. 1), and maps several reads that other do not (see Supplementary Data). It also can map read with fewer errors, and find more *loci* per read. Time-wise, srnaMapper is slower than methods that map significantly less reads, but comparable with the tools that map almost the same number of reads.

**Figure 1:**
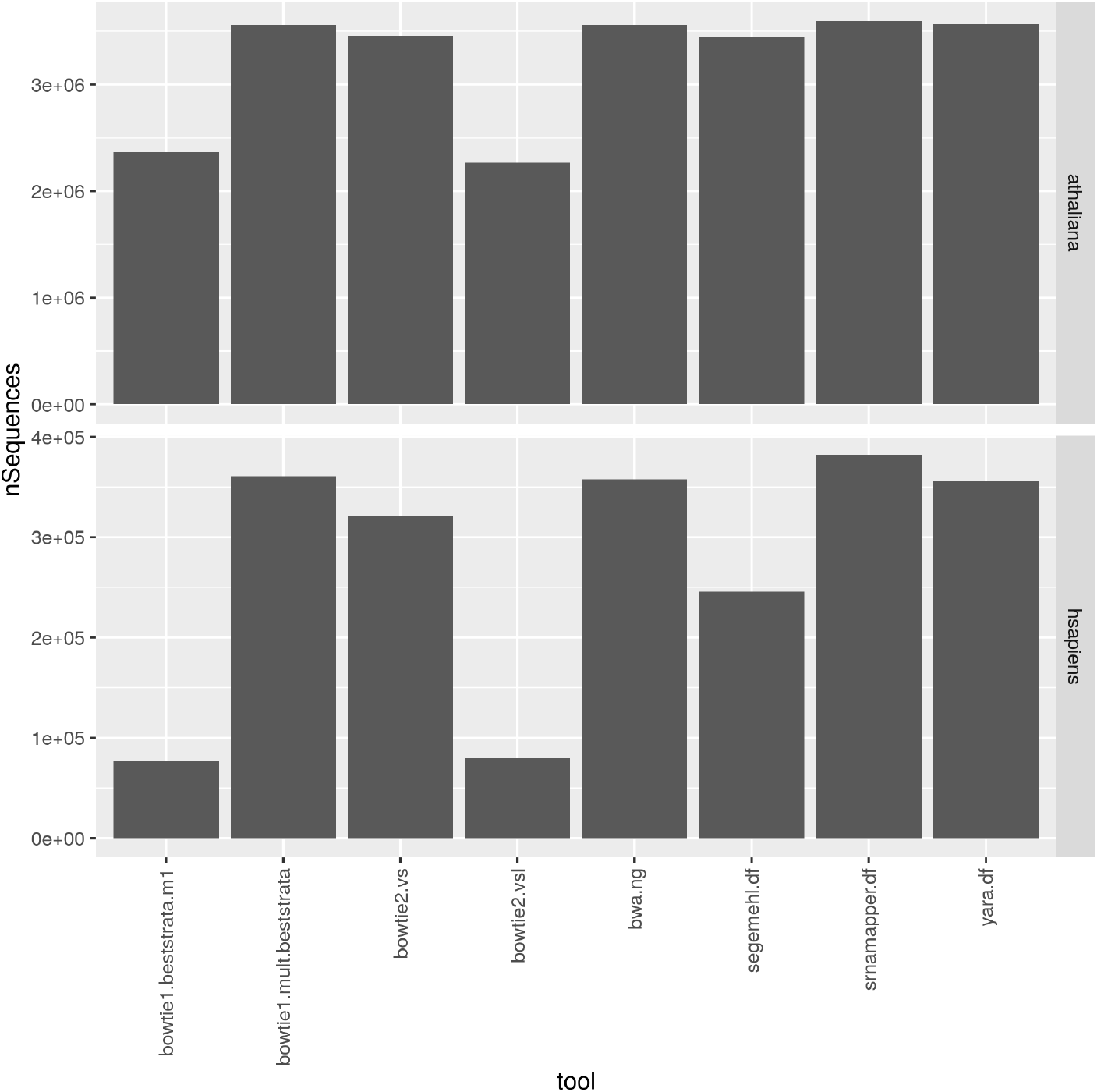
Number of reads mapped.

**Figure 2:**
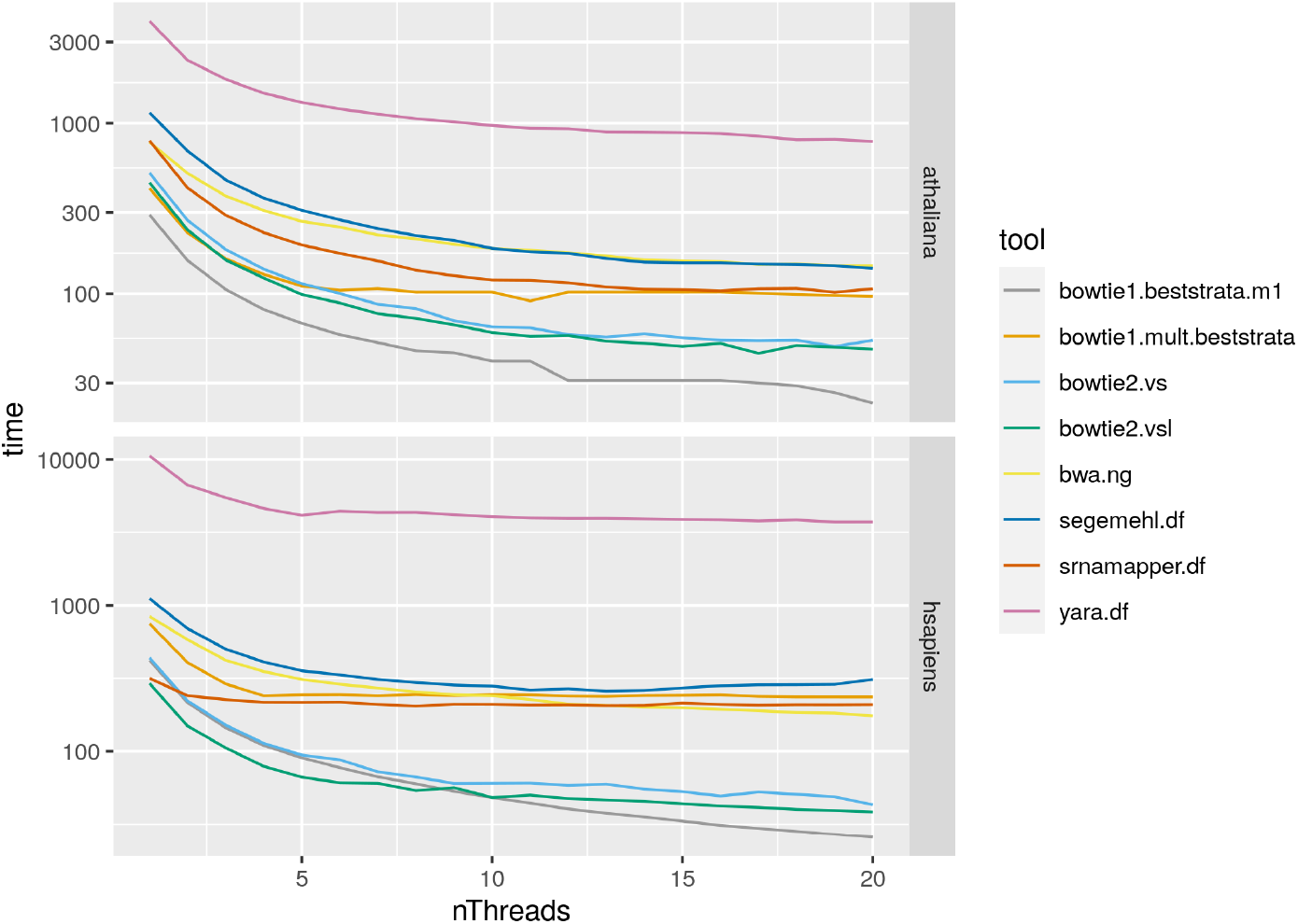
Time (in seconds) needed to map the reads. The y-axis is in log scale.

We believe that srnaMapper could be the tool of choice for mapping short-RNA reads, since it maps more reads, at more locations, with a modest time difference when compared to other best tools.

