## Supplementary data for "srnaMapper: an optimal mapping tool for sRNA-Seq reads"

#### 1 Information about the reads

The *Arabidopsis thaliana* dataset was first published in [1], and the *Homo sapiens*, in [2]. The datasets were downloaded from SRA, and processed as follows.

```
fasterq-dump -e 6 -p file.sra
fastx_clipper -Q33 -a adapter -l 15 -i file.fastq > file_trim.fastq
```

The number of reads per sample is given in Figure 1.
The distribution of sizes Figure 2.

#### 2 Information about the tools

Tool versions:

- bowtie2: 2.4.1
- bowtie: 1.3.0
- bwa: 0.7.17
- segemehl: 0.3.4
- yara: 0.9.11

Tool commands:

- srnamapper.df:
  ```
  srnaMapper -t <THREADS> -r <READS> -g <GENOME> -o <SAM> -n 100 -f 100
  ```
- Bowtie2.vsl:
  ```
  bowtie2 --very-sensitive-local -p <THREADS> -x <GENOME> -U <READS> -S <SAM>
  ```
- Bowtie2.vs:
  ```
  bowtie2 --very-sensitive -p <THREADS> -x <GENOME> -U <READS> -S <SAM>
  ```
- Bowtie1.beststrata.m1:
  ```
  bowtie --best --strata -k 1 -m 1 -S -p <THREADS> -x <GENOME> <READS> <SAM>
  ```
- BWA.ng:
  ```
  bwa aln -o 0 -t <THREADS> -f tmp.sai <GENOME> <READS> && bwa samse -f <SAM> <GENOME>.fa tmp.sai <READS>
  ```

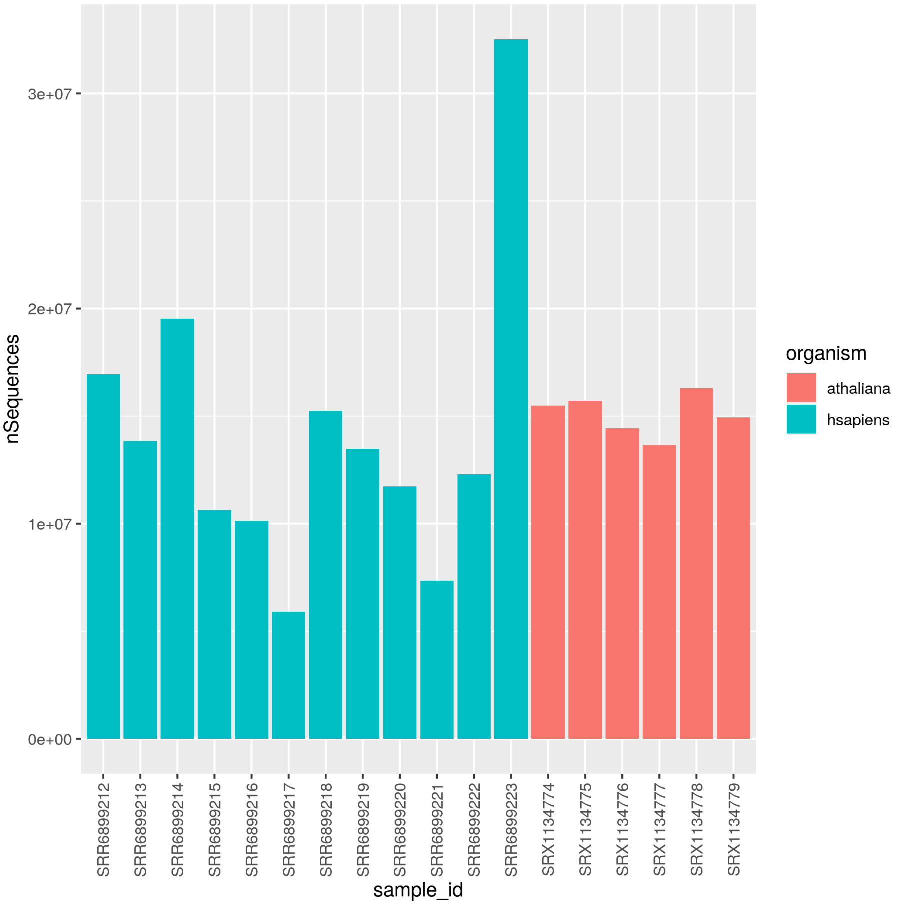


Figure 1: Number of reads per dataset.

- Bowtie1.mult.beststrata: `bowtie -k 100 --best --strata -S -p <THREADS> -x <GENOME> <READS> <SAM>`
- segemehl.df: `segemehl.x -t <THREADS> -d <GENOME> -i <GENOME>.idx -q <READS> -o <SAM>`
- yara.df: `yara_mapper --version-check FALSE -o <SAM> -e 10 -s 0 -y full -sa tag -t <THREADS> <GENOME> <READS>`

### 3 Time spent by srnaMapper with different files

We used srnaMapper with several FASTQ files. All the sequences of the different files are compiled into a unique reads tree, which is then mapped to the

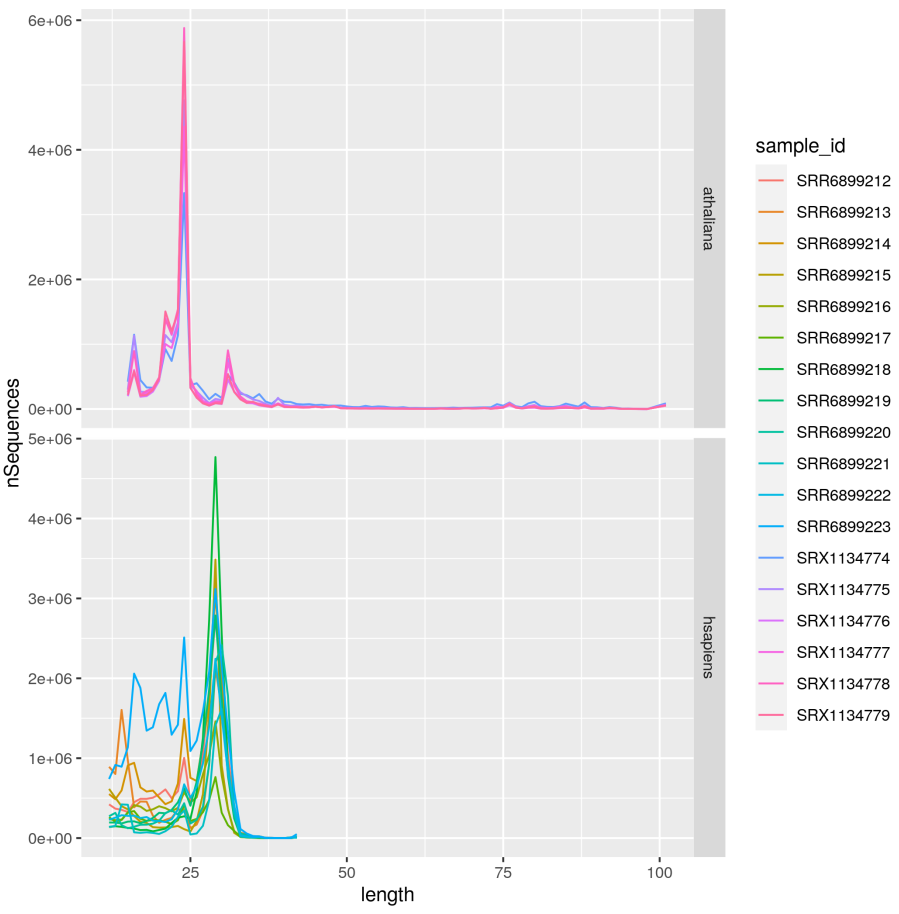


Figure 2: Distribution of the sizes of the reads.

genome. Results show that the time complexity is sub-linear (cf. Figure 3), which confirms that the approach is valid.

### 4 Comparison between srnaMapper and the other tools

The number of reads mapped by srnaMapper, and not the other tools, is given in Figure 5. Note that some reads (very few, not visible in the graph) are not mapped by srnaMapper, but are discarded because they are highly repeated (>500 times).

Sometimes, a read can be mapped by both srnaMapper and another tool. However, srnaMapper can map it with fewer errors, because, contrary to other

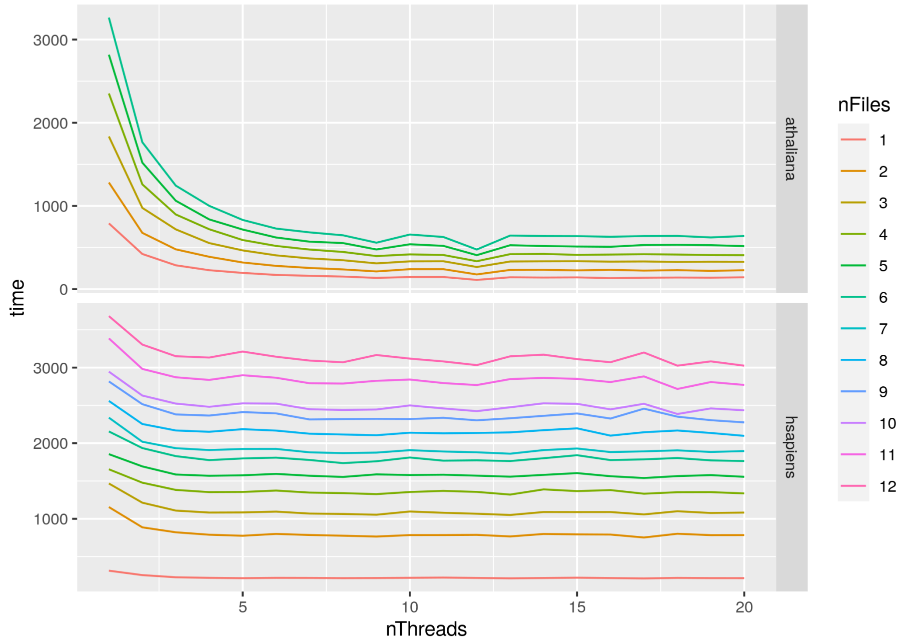


Figure 3: Time spent by srnaMapper, used with several files.

tools, srnaMapper do not resort to heuristics to map reads. The number of such reads is given in Figure 5

Last, a read can be mapped with the same number of errors, but srnaMapper can find more possible hits. The number of such reads is given in Figure **??**

#### 5 Detection of read edition

The SAM file produced by the mapped can help understanding where sequence edition takes place. For each mapping tool, we counted the number of substitutions, deletion, or insertions, and classified them in 5' edition if they are located at the 5' end, 3' edition, or interior edition otherwise. Notice that Yara do not indicate the substitutions the CIGAR format, and do not fill the MD tag. Yara can thus not be used for finding possible editions, and have been excluded from this analysis. The number of editions found is provided in Figure 7. These results confirm that 3' edition is slightly more frequent than 5' edition.

The previous previous graph shows, that in some cases, other tools find more edition than srnaMapper, which seemingly contradict previous results. The reason is that some of the edited reads map several times. Mapping tools which choose a random location may suggest an edition, which may or may not be true. When tools reported several hits, we classified as "ambiguous" editions which are not consistent for every hit. The number of ambiguous editions is

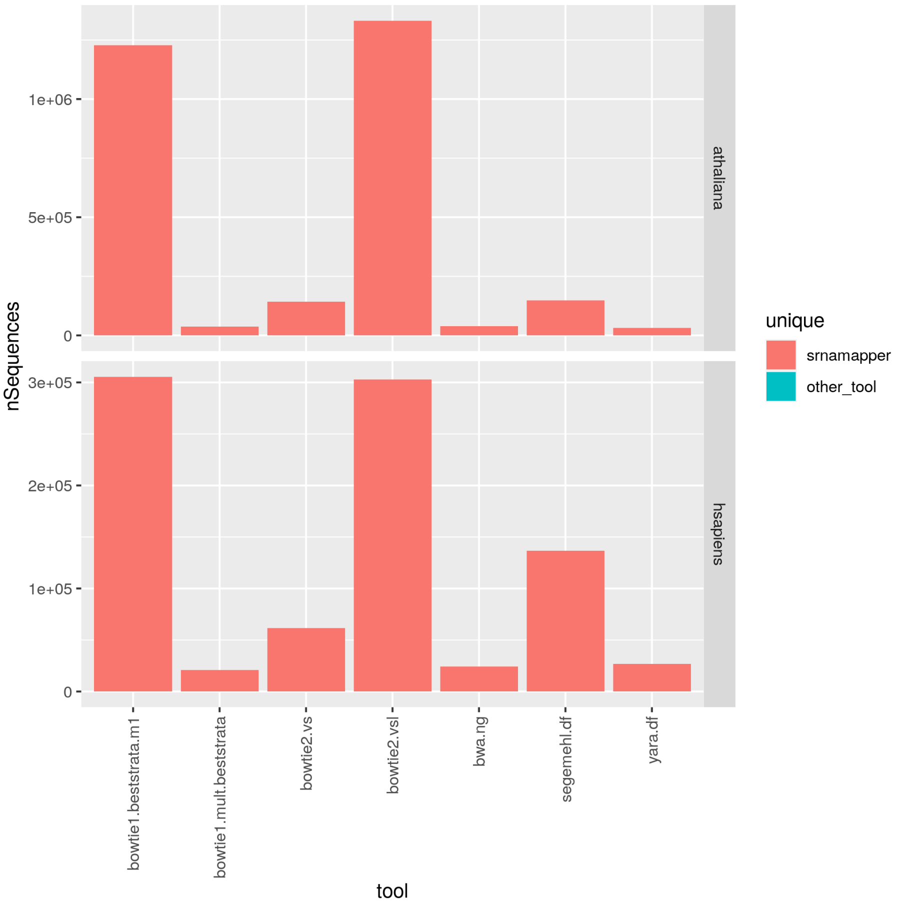


Figure 4: Number of reads mapped by srnaMapper, and not the other tools.

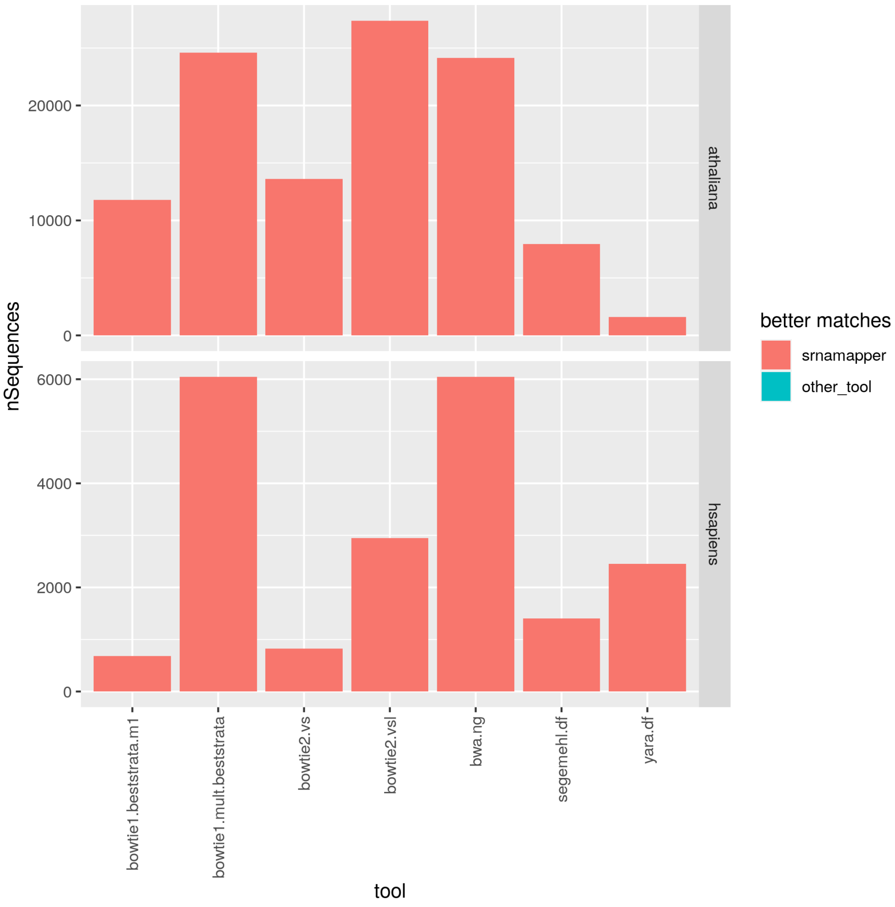


Figure 5: Number of reads mapped with fewer errors with srnaMapper.

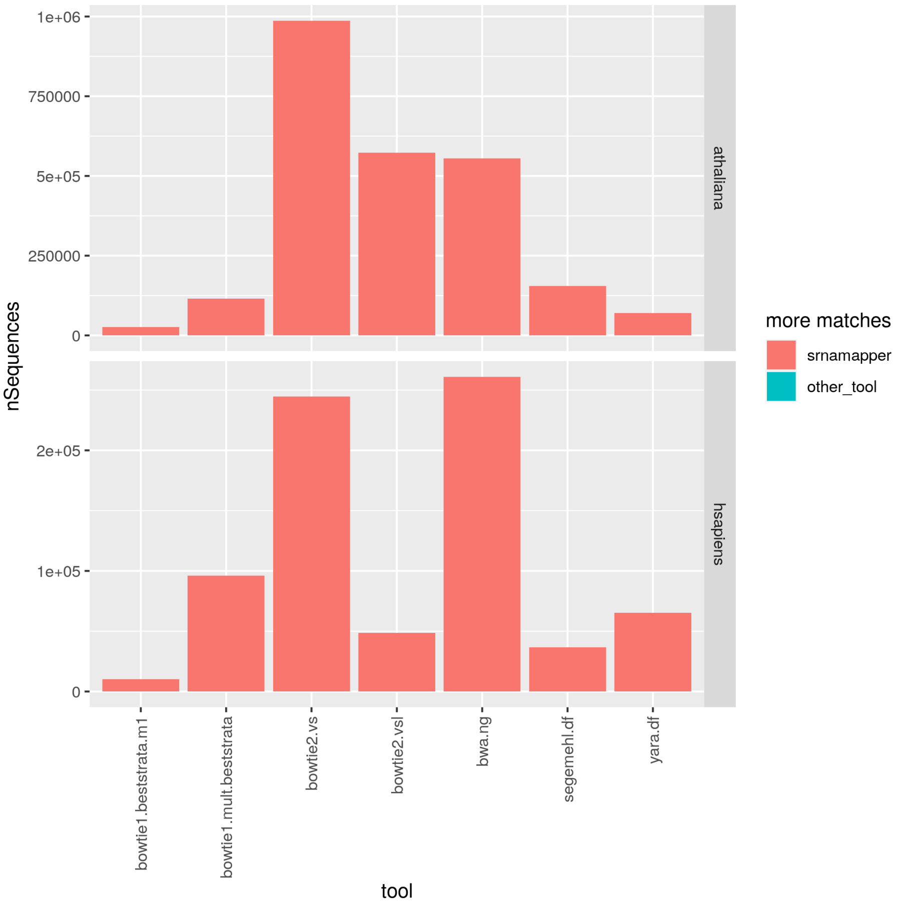


Figure 6: Number of reads mapped on more *loci* with srnaMapper.

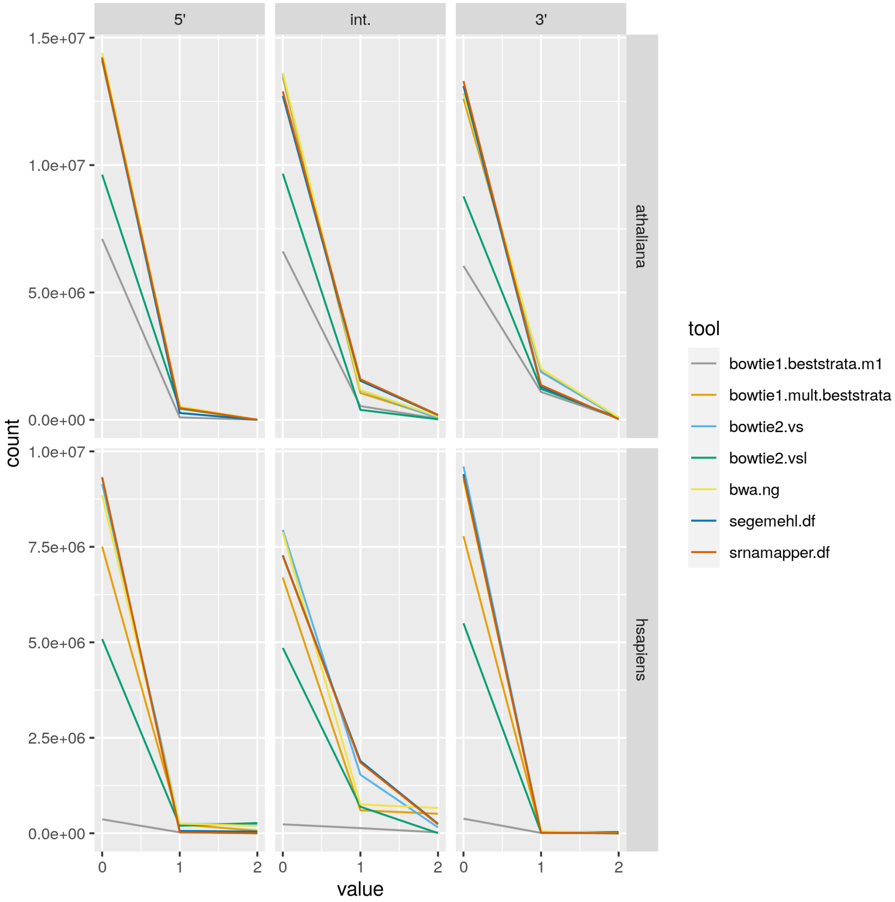


Figure 7: Number of reads mapped on more *loci* with srnaMapper.

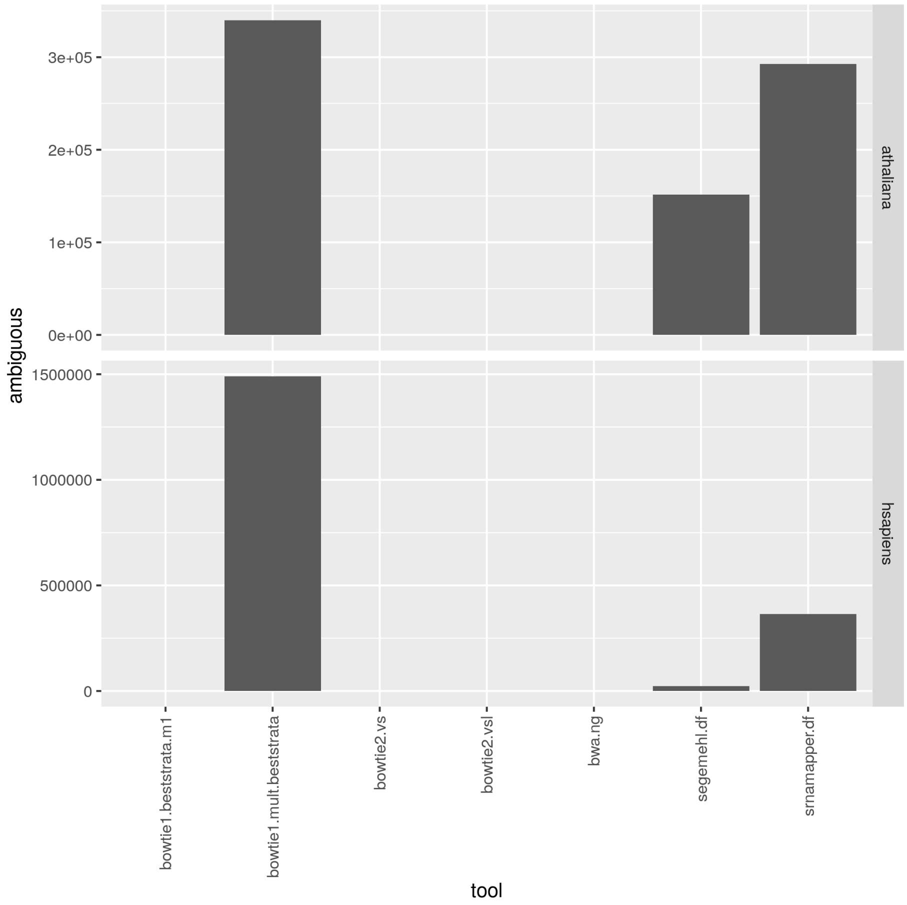


Figure 8: Number of ambiguous editions.

given in Figure 8.
